# LinAliFold and CentroidLinAliFold: Fast RNA consensus secondary structure prediction for aligned sequences using beam search methods

**DOI:** 10.1101/2022.06.17.496559

**Authors:** Tsukasa Fukunaga, Michiaki Hamada

## Abstract

RNA consensus secondary structure prediction from aligned sequences is a powerful approach for improving the secondary structure prediction accuracy. However, because the computational complexities of conventional prediction tools scale with the cube of the alignment lengths, their application to long RNA sequences, such as viral RNAs or long non-coding RNAs, requires significant computational time. In this study, we developed LinAliFold and CentroidLinAliFold, fast RNA consensus secondary structure prediction tools based on minimum free energy and maximum expected accuracy principles, respectively. We achieved software acceleration using beam search methods that were successfully used for fast secondary structure prediction from a single RNA sequence. Benchmark analyses showed that LinAliFold and CentroidLinAliFold were much faster than the existing methods while preserving the prediction accuracy. As an empirical application, we predicted the consensus secondary structure of coronaviruses with approximately 30,000 nt in 5 and 76 minutes by LinAliFold and CentroidLinAliFold, respectively. We confirmed that the predicted consensus secondary structure of coronaviruses was consistent with the experimental results. The source code is freely available at https://github.com/fukunagatsu/LinAliFold-CentroidLinAliFold.

## Introduction

RNAs play essential roles in various biological processes, such as transcriptional regulation and nucleotide modification ([1, 2]). RNA function is closely related to their structures, and as a result many studies have been conducted to determine RNA structures experimentally ([3]). However, these experiments are too expensive and time-consuming to perform the large-scale RNA structure analysis; therefore, a fast computational structure prediction analysis is widely used as an alternative approach. Although protein tertiary structures can be predicted with high accuracy using AlphaFold2 ([4]), the prediction of RNA tertiary structures with acceptable performance remains a challenge ([5]). Accordingly, RNA secondary structure prediction analyses are frequently performed, and various secondary structure prediction tools have been developed ([6, 7, 8, 9, 10, 11]).

RNA consensus secondary structure prediction from sequence alignments is a powerful approach for accurate secondary structure prediction ([12, 13, 14, 15, 16]). These methods improve prediction performance by taking advantage of the fact that evolutionarily conserved base-pairs tend to form secondary structures ([17]). The conventional method is RNAalifold, which predicts a consensus minimum free energy (MFE) structure while including sequence covariation scores in the energy model ([12, 14]). Another representative tool is CentroidAlifold, which computes the centroid structure in the probability distribution of candidate structures based on the maximum expected accuracy (MEA) principle ([15]).

These prediction methods are highly accurate and are widely used by experimental biologists. However, they are too computationally expensive to be applied to long sequences, such as viral RNAs and long non-coding RNAs, because the computational complexities are proportional to the cube of the alignment lengths. One solution to this problem is the local folding method, which reduces the computational time by ignoring long-range base-pair interactions ([18, 19, 20, 21]). This approach enables analysis of RNA secondary structures at the transcriptome level ([22, 23]). However, ignored long-range interactions are often found in natural RNAs, some of which have functional roles such as regulation of splicing or transcription ([24, 25]). For example, the SARS-CoV-2 RNA genome forms 29.8knt spanning long-range base-pairs that are presumed to contribute to genome replication ([26]).

As another approach for long RNA prediction, LinearFold was recently proposed by Huang *et al*. for fast secondary structure prediction from a single RNA sequence. LinearFold accelerates structure prediction by pruning candidate structures using a beam search method ([27]). In other words, by using heuristics to obtain approximate solutions, LinearFold can predict secondary structures including long-range interactions, within a reasonable amount of time. Despite the use of the heuristics, Huang *et al*. demonstrated the superior prediction accuracy of LinearFold over the conventional prediction tools, RNAfold and CONTRAFold. The beam search-based acceleration technique is currently applied to various RNA secondary structure analyses, including partition function calculation ([28]), structural alignment ([29]), stochastic sampling ([30]), sequence design ([31]), and pseudoknot prediction ([32, 33]).

In the current study, we developed fast RNA consensus secondary structure prediction tools using beam search methods. We implemented two prediction tools, LinAliFold and CentroidLinAliFold, accelerated methods of RNAalifold and CentroidAlifold by beam search methods, respectively. We confirmed that LinAliFold and CentroidLinAliFold showed comparable prediction accuracy to existing programs and that the two programs were much faster than existing programs.

## Methods

### The LinAliFold algorithm

We first briefly review the RNAalifold algorithm, which outputs a consensus MFE structure from the input alignment data under a thermodynamic energy model incorporating sequence covariation scores ([12, 14]). The prediction algorithm is based on dynamic programming (DP), with time and space complexities of *O*(*S N*^3^) and *O*(*N*^2^) for a sequence alignment whose the number of sequences is *S* and the length is *N*. As a thermodynamic energy model, RNAalifold uses the same neighborhood energy model as RNAfold, which predicts the MFE structure from a single sequence. RNAalifold is unique as it uses sequence covariation scores for structure prediction.

The sequence covariation score is defined for two alignment columns, and the columns with higher scores are less likely to become base-paired in the predicted structure. The score is designed to have a lower value when there are more base mutations such that base pairing is conserved in the two columns. Conversely, when gaps or two bases that do not form a base-pair are included in the two columns, the value is designed to be higher. The sequence covariation score model was firstly designed in RNAalifold, and this score model regarded all substitution rates between base pairs as equal. We refer to this sequence covariation score model as the simple score model. RNAalifold also provides the RIBOSUM score model, which reflects substitution rates between base pairs in real data. Specifically, the model was learned from an alignment of 2492 SSU ribosomal RNAs from the European Ribosomal RNA Database ([34, 35, 14]).

We also review the beam search method in RNA secondary structure analysis ([27]). As an example, consider evaluating the energy score when a column *i* forms a base-pair with a column on the 5^′^ side of the column *i*. In conventional exact approaches, the energy scores of all columns that can form base-pairs are investigated, and all energy scores are retained for subsequent calculations in the DP. In contrast, the heuristic beam search method evaluates only the energy scores of columns whose scores are retained. Subsequently, the method only maintains the energy scores of the top *b* columns with the lowest energy scores, and discards the energy scores of the other columns. Here, before the column *i* is evaluated, all columns on the 5^′^ side from the column *i* must have been evaluated. Because this condition is not satisfied by a bottom-up DP, which is conventionally used in RNA structural analysis, LinearFold employed a left-to-right DP instead ([36, 37]). This beam search heuristics was also applied to the evaluation of multi loop formation. LinearFold achieved significant acceleration by pruning the search space with this beam search, and reduced the time and space complexities from *O*(*N*^3^) and *O*(*N*^2^) to *O*(*Nb*log*b*) and *O*(*Nb*). *b* is the beam size and is a user-defined parameter.

LinAliFold is a program that speeds up RNAalifold by using the beam search method. Specifically, the bottom-up DP in RNAalifold was replaced by the leftto-right DP, and the beam pruning was applied to the DP calculation. Compared with RNAalifold, the time complexity is reduced from *O*(*S N*^3^) to *O*(*S Nb*log*b*). On the other hand, the space complexity is the same *O*(*N*^2^) as RNAalifold because we used two-dimensional arrays for the data structure to store the sequence covariance scores. Note that we can reduce the spatial complexity by using hashes instead of two-dimensional arrays. However, we did not use hashes because preliminary experiments showed that the usage of hashes worsened the computational speed performance. In addition, we have slightly modified the calculation method of RNAalifold and LinearFold; this is described in the Supplementary Material.

### The CentroidLinAliFold algorithm

We next present an overview of the CentroidAlifold algorithm ([15]). CentroidAlifold was designed from the viewpoint of MEA, which maximizes the expected prediction accuracy under a probability distribution on candidate solutions. In particular, CentroidAlifold utilizes the γ-centroid estimator ([7]), which is theoretically superior to the frequently used ContraFold MEA estimator ([6]).

CentroidAlifold first computes 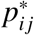, a base pairing probability (BPP) between two columns *i* and *j*, for all column pairs, as follows:

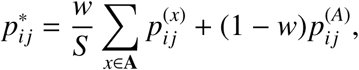

where *w* ∈ [0, 1] is a mixture weight parameter, 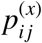 is the BPP on an individual sequence *x*, and 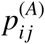 is the BPP on the input alignment **A**. Here, 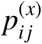 and 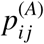 can be computed using the McCaskill algorithm and the extension of RNAalifold with the time complexities *O*(*N*^3^) and *O*(*S N*^3^), respectively.

CentroidAlifold then performed Nussinov-style DP using 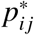 as follows ([38]):

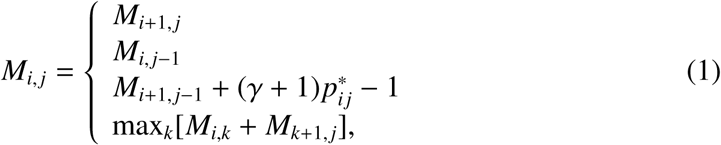

where *M*_*i, j*_ denotes the best score for **A**[*i*.. *j*], which is defined as a subsequence between *i* and *j* on alignment **A**.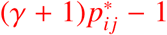 in the third line of equation (1) represents that two columns with higher BPP are more likely to be included in the predicted secondary structure. γ is a user-defined hyperparameter that controls the number of base pairs in the predicted structure. A higher γ value is expected to increase the number of base pairs, i.e., increase the sensitivity (SEN) and decrease the positive prediction value (PPV). The time and space complexities of DP are *O*(*N*^3^) and *O*(*N*^2^). CentroidAliFold finally predicted the secondary structure by traceback of the DP matrix. Accordingly, the total time and space complexities of CentroidAlifold were *O*(*S N*^3^) and *O*(*N*^2^).

CentroidLinAliFold is the software that speeds up CentroidAlifold by applying the beam search method to each step of CentroidAlifold. The calculation method of 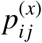 using beam search has been proposed as LinearPartition ([28]) with a time complexity *O*(*Nb*^2^), and the algorithm was reimplemented and incorporated into CentroidLinAliFold. In addition, to calculating 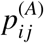, we developed a novel beam search-based algorithm with time complexity *O*(*S Nb*^2^) by extending LinAliFold. The beam search method was not applied to the Nussinov-style DP step because this step empirically requires less computational time when using the threshold method. The threshold method is a fast structure prediction method that uses only 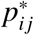 larger than the threshold in DP recursion. From the third line of equation (1), we can set the threshold to 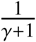 without affecting the prediction results. Therefore, the number of base pairs to be considered per sequence position is bounded by *O*(γ), and the time complexity of this step becomes *O*(γ*N*^2^).

### Datasets and evaluation measures

We evaluated the prediction performance using eight RNA families provided in the RNAStralign dataset: 5S rRNA, 16S rRNA, group I intron, RNase P, SRP RNA, telomerase RNA, tmRNA, and tRNA ([39]). We randomly selected 5, 10, or 20 sequences for each RNA family, and aligned the sequences by mafft-xinsi v7.490, which performs multiple alignments considering RNA secondary structure information ([40, 41]). Note that most current RNA structural alignment tools require at least the cube of the sequence length in the computation time, so the advantage of the beam search-based consensus structure prediction method may appear to have diminished. However, because we can develop fast structural alignment tools using the beam search methods, this drawback will be disappeared in the future.

We then predicted the consensus secondary structures from the alignments, and evaluated the prediction quality of individual RNA sequences using three metrics: PPV, SEN, and the Matthews correlation coefficient (MCC) with respect to base-pairs ([15]). We prepared five datasets for each evaluation experiment, and measured the average values of the evaluation metrics.

The maximum alignment length of the RNAStralign dataset is approximately 1,500 bp for 16S rRNA, and the alignment length is insufficient to evaluate the computation speed of LinAliFold and CentroidLinAliFold. To test the performance on longer sequences, we created artificial alignment datasets by concatenating multiple alignments of 16S rRNAs two to ten times. For each number of concatenations, we evaluated the average computational time and required memory size for the five datasets. Although the number of concatenations is the same, the alignment lengths differed for each dataset. Therefore, we regarded the average alignment lengths as the representative alignment length for the number of concatenations. In this experiment, the computation was conducted on an Intel Core i5 2.4 GHz CPU with 8GB of memory.

For practical application of long RNA sequences, we predicted consensus secondary structures of coronaviruses with sequence lengths of approximately 30,000 nt. We used the coronavirus genome dataset of 25 sequences compiled by Li *et al*. ([29]). The dataset consisted of 16 SARS-CoV-2, five human SARS-CoV-1, and four bat coronavirus genomes. We could not apply mafft-xinsi to the coronavirus dataset owing to the shortage of memory;therefore, we used mafft with default settings for multiple sequence alignments. An experimentally determinedmreference RNA secondary structure, we used the structure determined by Ziv *et al*., including long-range RNA-RNA interactions ([26]). The computation was performed on an Intel Xeon Gold 6154 4.0 GHz CPU with 128GB memory in this experiment.

We compared the performance of LinAliFold and CentroidLinAliFold with those of RNAalifold and CentroidAlifold, respectively. We also used RNAfold and Centroidfold to predict the secondary structure from a single sequence to compare the prediction performance. For the thermodynamic energy parameter, we used Andronescu’s BL* parameter owing to its high prediction accuracy ([42]). As in previous studies, we used 100 and 0.5 as the default values of the beam size *b* and the mixture weight parameter *w*, respectively ([15, 27]). We also used the simple score model as the default sequence covariation score model.

## Results

### Evaluation of prediction accuracy

We first evaluated the prediction accuracy of the six RNA structure prediction tools using the RNAStralign dataset. Fig. 1 and Fig. S1 show the average prediction results for all RNA families. MEA-based tools have multiple prediction results depending on γ, and we selected γ from 2^*k*^ : 1 ≤ *k* ≤ 7, *k* N. We found that the consensus secondary structure prediction tools were superior to the prediction tools from single sequences, and that the MEA-based tools performed better than the MFE-based tools. These results are consistent with previous studies. In addition, we verified that LinAliFold and CentroidLinAliFold have comparable prediction accuracy to RNAalifold and CentroidAlifold, respectively. These results were independent of the number of sequences in the alignments (Fig. S1). Table 1 and Table S1-8 show the prediction performance of each RNA family. In this analysis, the hyperparameter γ was automatically determined based on the maximization of the pseudo-expected accuracy of the MCC ([43]). The pseudo-expected accuracy of the MCC is an approximate MCC estimator, and it is computed from the BPPs only, without using a reference structure. We found that the consensus secondary structure prediction tools performed well. However, the prediction from a single sequence was more accurate for some families, such as group I introns and tmRNAs. This result may be due to inaccurate structural alignments. Additionally, we confirmed that LinAliFold and CentroidLinAliFold have equal or better accuracy than RNAalifold and CentroidAlifold, respectively. In particular, CentroidLinAliFold achieved the best prediction performance with an MCC of 0.702 when the the number of sequences was 20. These results suggest that the beam search heuristics do not have a negative influence on the prediction accuracy of RNA consensus secondary structure prediction.

**Table 1:** The MCC results of prediction accuracy for each RNA family when the number of sequences was 20

| RNA Family | RNAfold | CentroidFold | RNAalifold | LinAliFold | CentroidAlifold | CentroidLinAliFold |
| --- | --- | --- | --- | --- | --- | --- |
| 5s rRNA | 0.729 | 0.676 | 0.885 | 0.898 | 0.885 | **0.903** |
| 16S rRNA | 0.434 | 0.520 | 0.684 | 0.656 | **0.734** | 0.720 |
| group I intron | 0.708 | **0.752** | 0.598 | 0.632 | 0.627 | 0.644 |
| RNaseP | 0.568 | 0.647 | 0.616 | 0.596 | 0.635 | **0.654** |
| SRP RNA | 0.643 | 0.640 | 0.596 | 0.645 | 0.602 | **0.649** |
| telomerase RNA | 0.455 | 0.493 | 0.436 | 0.555 | 0.551 | **0.572** |
| tmRNA | 0.512 | **0.547** | 0.382 | 0.475 | 0.460 | 0.505 |
| tRNA | 0.809 | 0.784 | 0.962 | **0.966** | 0.962 | **0.966** |
| average | 0.607 | 0.632 | 0.645 | 0.678 | 0.682 | **0.702** |

The bold values are the highest scores in each row.

**Figure 1:**
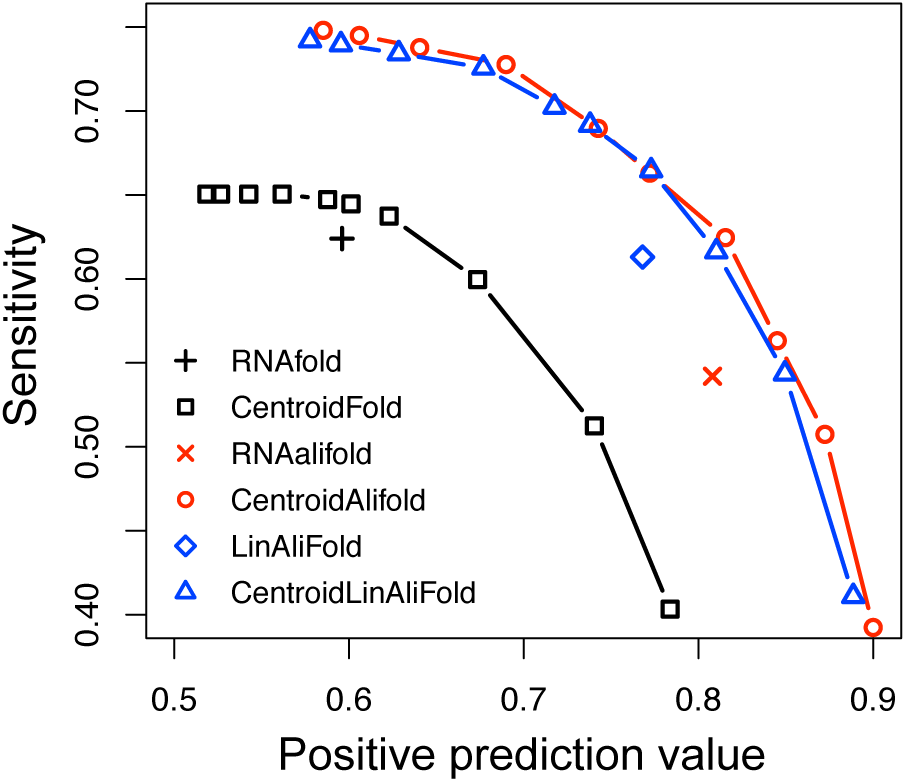
Prediction performance of RNA secondary structure on the RNAStralign dataset with 20 sequences. The x-axis and the y-axis represent PPV and SEN, respectively. Symbols and colors used above represent: RNAfold (black crosses), CentroidFold (black squares), RNAalifold (red crosses), CentroidAlifold (red circles), LinAliFold (blue squares), and CentroidLinAliFold (blue triangles).

### Evaluation of computational time and required memory size

We next assessed the computational time required by the developed software (Fig. 2 and Fig. S2). We confirmed that LinAliFold and CentroidLinAliFold could perform the prediction in almost linear computational time with respect to the alignment lengths. In addition, LinAliFold and CentroidLinAliFold were considerably faster than RNAalifold and CentroidAlifold when the alignment lengths were long, and the speed efficiency increased as the alignment lengths increased. For example, with an alignment length of 15,600 nt and the number of sequences of 20, LinAliFold was 7.8 times faster than RNAalifold. As another example, with an alignment length of 4,680 nt and the number of sequences of 20, CentroidLinAliFold was 20.9 times faster than CentroidAlifold. Furthermore, LinAliFold was found to be largely faster than CentroidLinAliFold. There are various reasons for this difference. First, the computational complexity is different; LinAliFold is *O*(*S Nblogb*), whereas CentroidLinAliFold is *O*(*S Nb*^2^). Second, LinAliFold only needs to perform DP once for an alignment, like the Viterbi algorithm of hidden Markov models (HMMs), whereas CentroidLinAliFold needs to perform DP twice for a sequence or alignment, like the forward-backward algorithm of HMMs. Third, CentroidLinAliFold has to calculate BPPs for individual sequences as well as for alignments. Fourth, CentroidAliFold has to calculate the BPPs in the log space to avoid underflow, but LinAliFold does not use the log space.

**Figure 2:**
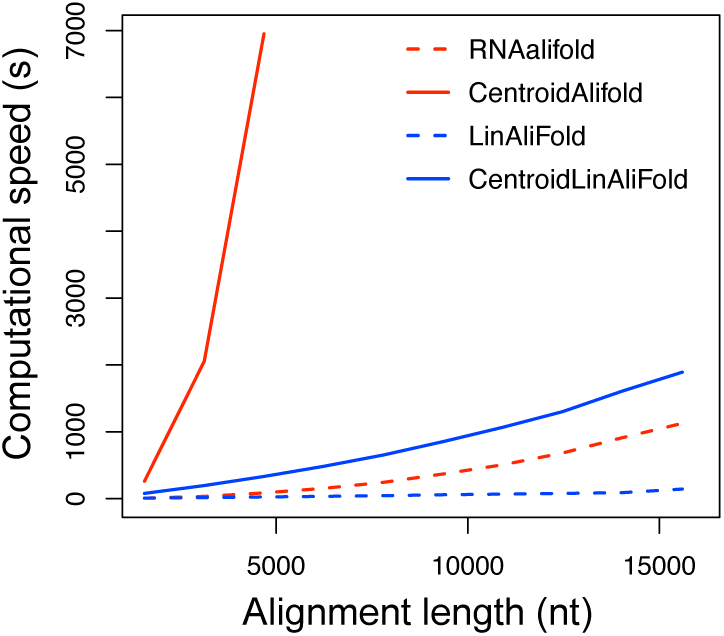
Evaluation of the computational time when the number of sequence was 20. The xand y-axis represent the alignment lengths (nt) and computational time (s), respectively. The lines represent RNAalifold (red dashed), CentroidAlifold (red solid), LinAliFold (blue dashed), and CentroidLinAliFold (blue solid). The exact computational time was not measured when it exceeded 2 hours.

We also investigated the required memory size for the developed software (Fig. 3 and Fig. S3). The results show that RNAalifold and LinAliFold required almost the same amount of memory size, but CentroidLinAliFold was more memory efficient than CentroidAlifold. Because CentroidAlifold and CentroidLinAliFold should have equivalent amounts of memory based on the algorithm, this difference in results may be due to implementation rather than the algorithms. In addition, we verified that LinAliFold was better memory efficiency than CentroidLinAliFold. The main reason for this difference is that calculating BPPs requires more internal variables than finding an MFE structure. We further examined the dependencies of computational time and required memory size on the number of sequences (Tables S9-10). The computational time increased for both LinAliFold and CentroidLinAliFold as the the number of sequences increased; however, the required memory size did not change significantly.

**Figure 3:**
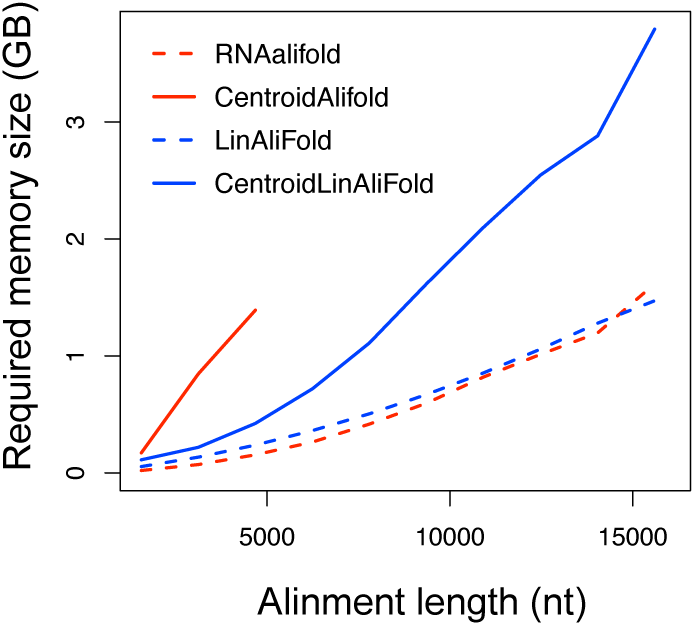
The required memory size was evaluated when the number of sequences was 20. The xand y-axis represent the alignment lengths (nt) and required memory size (GB), respectively. The lines represent RNAalifold (red dashed), CentroidAlifold (red solid), LinAliFold (blue dashed), and CentroidLinAliFold (blue solid). The required memory size was not measured when computational time exceeded 2 hours.

### Analysis of the dependence of the performance on the sequence covariation score model and the beam size

We then analyzed the effects of the sequence covariation score model on the prediction accuracy (Table S11). Note that we could not perform the analysis for CentroidAlifold because this tool does not have the option of changing the sequence covariation score model. As expected from the results of a previous study ([14]), the RIBOSUM model outperforms the simple score model with any prediction tool. When comparing the prediction performance for each RNA family using the RIBOSUM model, some families improved the prediction accuracy significantly, and no families showed substantial decreases.

We also estimated the influence of the beam size *b* on the performance of LinAliFold and CentroidLinAliFold. Tables S12-13 show the MCC changes with beam size *b*. The prediction accuracy monotonically increased with increasing *b* as expected, but the performance did not improve significantly over *b* = 100. Tables S14-15 show the changes in the computational time and required memory size with the change in beam size *b*. We found that the computational time increased but the required memory size did not change significantly as *b* increased. These results are consistent with the results of the computational complexity analysis.

### Prediction of coronavirus RNA consensus secondary structure

We finally predicted the consensus secondary structures of coronaviruses using LinAlifold and CentroidLinAliFold. Table 2 shows the computational time and required memory size for the prediction. The computational times include the execution time of MAFFT for our tools; however, most of the required computational time is the structure prediction because MAFFT completed the computation in seven seconds. We demonstrated that our tools could predict the secondary structure of coronaviruses within a reasonable amount of time and memory. Note that LinearTurboFold, a beam search-based structural alignment and structure prediction tool ([29]), required 18h11m1s time and 44.1 GB memory for coronavirus secondary structure prediction. This difference in software performance between our tools and LinearTurboFold may be attributed to differences in the purpose of programs; LinearTurboFold performs structural alignment, but our tools do not perform it.

**Table 2:** The computational time and required memory size for coronavirus secondary structure prediction

| software | computational time | required memory |
| --- | --- | --- |
| LinAliFold | 4m48s | 4.7G |
| CentroidLinAliFold | 1h15m24s | 14.9G |
| LinearTurboFold | 18h11m1s | 44.1G |

We compared the predicted and experimentally determined structures of canonical 5^′^-UTRs, which have been well studied. We confirmed that LinAliFold and CentroidLinAliFold produce predictions consistent with the experimental results, with a few exceptions, such as the shortage in the stem structure of the SL6 motif and the lack of the SL5-3 motif, respectively (Fig. 4 and S4). A previous study showed that the beam search-based prediction from a single sequence could not predict the consistent structures with experiments ([29]). Therefore, prediction results by LinAliFold and CentroidLinAliFold demonstrated that RNA consensus secondary structure prediction using beam search is an effective approach for viral RNA structure prediction.

**Figure 4:**
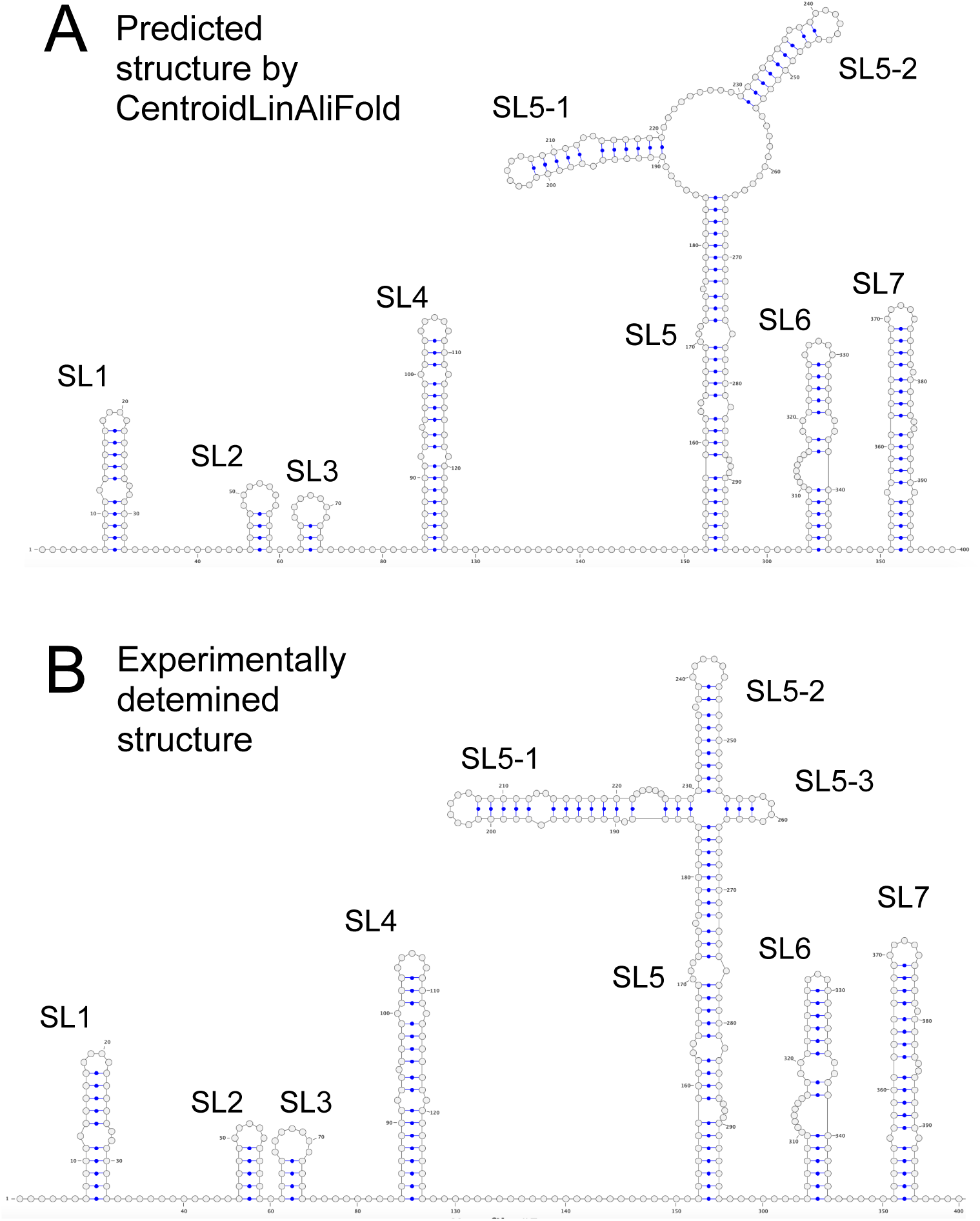
(A) A predicted RNA consensus secondary structure of coronavirus 5^′^-UTR by CentroidLinAliFold. (B) An experimentally determined secondary structure of the coronavirus. Graphical views were created using VARNA ([49]).

The region forming the SL3 motif in the 5^′^-UTR of the coronavirus can form an alternative structure that takes a long-range interaction spanning 29.8 knt.

However, most RNA secondary structure prediction tools predict only one of two different structures. For example, while LinearTurboFold predicted a structure with the long-range interaction, our tools predicted the canonical structure. However, prediction methods that calculate BPPs, such as LinearTurboFold and CentroidAliFold, can estimate the possibility that alternative structures form. Therefore, we examined whether CentroidLinAliFold could capture the potential of the long-range interaction by examining BPPs. Specifically, we calculated the average probabilities that the bases constituting the SL3 motif formed the base pairs of each structure. As a result, we found that the probabilities that the region formed the canonical SL3 structure and the long-range interaction were 37% and 7%, respectively. This result indicates that CentroidLinAliFold can capture the long-range interactions in the coronavirus genome.

## Discussion

In this study, we developed LinAliFold and CentroidLinAliFold, which adopt beam search methods to accelerate RNA consensus secondary structure prediction. We investigated the prediction performance using the RNAStralign dataset, and verified that LinAliFold and CentroidLinAliFold have comparable accuracy to the existing programs, RNAalifold and CentroidAlifold. However, the computational speeds of LinAliFold and CentroidLinAliFold were much faster than those of the existing programs, especially for longer alignment lengths. In addition, we demonstrated that these two programs could accurately predict the secondary structures of coronaviruses of approximately 30,000 nt lengths in a reasonable amount of time.

While we were undergoing peer review for our paper, the paper about LinearAliFold, linear-time consensus secondary structure prediction tools based on the same concept as LinAliFold, was published in arxiv ([44]). Benchmark studies showed that LinearAliFold is also much faster than RNAalifold while preserving the prediction accuracy. Unlike the methods we developed, LinearAliFold can predict structures containing pseudoknots or sample the secondary structures from the stochastic distribution. On the other hand, unlike LinearAliFold, CentroidLinAliFold can predict sophisticated MEA structures based on the γ-centroid estimator. All of these methods will be foundational tools in predicting the common secondary structure of long RNAs.

We used the same score models for the sequence covariation score model as in the RNAalifold software was used. However, previous benchmarks suggested that the score models do not have high detection performance because the score models ignore the effects of stacking energy and nucleotide frequency distribution ([45], [46]). Rivas *et al*. recently showed that average product corrected (APC) G-test statistics have robust and sensitive performance for sequence covariation detection ([46]). This result suggests that the use of the APC G-test statistics may be effective for consensus secondary structure prediction. However, because the computational time of the APC G-test statistics scale with the square of the alignment lengths, the simple integration of the APC G-test statistics into LinAliFold or CentroidAlifold may result in the loss of the high speed advantage of these tools. We envision the development of a fast computable and sensitive sequence covariation score model.

The beam search method has been used to increase the speed of various RNA structure informatics tools, and the scope of its application should continue to expand. For example, CentroidHomfold ([47]) and CentroidAlign ([48]), which we previously developed for the structure prediction of a single sequence using homologous sequence information and structural alignment, can also be accelerated by beam search methods. The speedup of existing tools based on beam search methods is essential for genome-scale RNA secondary structural analysis in the research area of RNA informatics.

## Acknowledgments

Computations were partially performed on the NIG supercomputer at ROIS National Institute of Genetics and the SHIROKANE supercomputer at human genome center, the Institute of Medical Science, The University of Tokyo.

## Funding

This work was supported by the Japan Society for the Promotion of Science (grant numbers JP19K20395, JP20H05582 and JP22H04891 to T.F.).

## Notes

### Competing Interest Statement

The authors have declared no competing interest.

### Summary of Updates

We highlighted the changes in red color.

