## Supplementary Material for "LinAliFold and CentroidLinAliFold: Fast RNA consensus secondary structure prediction for aligned sequences using beam search methods"

#### Supplementary Methods

##### Further differences between the LinAliFold algorithm and previous tools

LinAliFold is an algorithm that applies a beam search method in LinearFold to RNAalifold, but LinAliFold has three other minor differences from RNAalifold and LinearFold.

The first is the calculation method of hairpin candidates. While LinearFold calculates $b$ hairpin candidates, LinAliFold calculates hairpin candidates with a loop length of $b$ bases or less. In other words, LinAliFold adopts a local folding method in the calculation of hairpin loops. This is because it was difficult to efficiently enumerate $b$ hairpin candidates in the RNA consensus secondary structure prediction. Note that this change should not significantly impact the prediction because hairpins with large loops are unlikely to occur.

The second is the energy evaluation of unpaired nucleotides in multi loops. While RNAalifold counts the numbers of nucleotides in individual sequences, LinAliFold approximates the number of nucleotides by the sub-alignment length in multi loops for fast computation. In other words, RNAaliflod does not include gaps on the multi loops in the numbers of unpaired nucleotides, but LinAliFold includes the gaps in the numbers.

The third is the evaluation of the gaps. While RNAalifold regards gaps as nearby bases when evaluating the type of bases, LinAliFold does not evaluate the gaps in such cases for ease of the implementation.

### Supplementary Figures

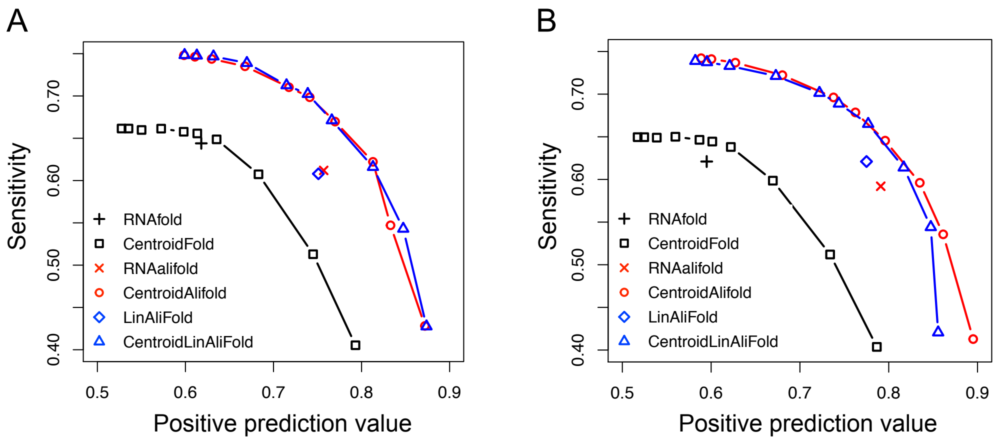


Fig. S1 Prediction performance of RNA secondary structure on the RNAStralign dataset with (A) 5 sequences and (B) 10 sequences. The x-axis and the y-axis represent PPV and SEN, respectively. Symbols and colors used above represent: RNAfold (black crosses), CentroidFold (black squares), RNAalifold (red crosses), CentroidAlifold (red circles), LinAliFold (blue squares), and CentroidLinAliFold (blue triangles).

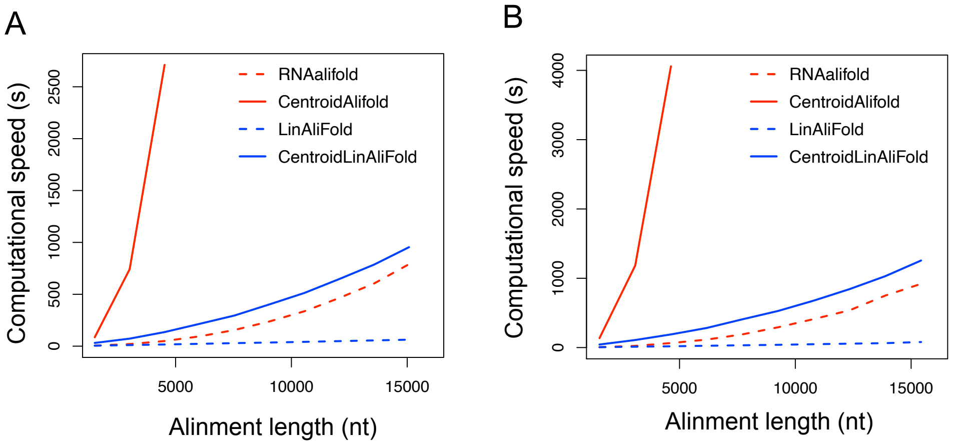


Fig. S2 Evaluation of the computational time for a sequence size of (A) 5 and (B) 10. The x- and y-axis represent the alignment lengths (nt) and computational time (s), respectively. The lines represent RNAalifold (red dashed), CentroidAlifold (red solid), LinAliFold (blue dashed), and CentroidLinAliFold (blue solid). The exact computational time was not measured when it exceeded 2 hours.

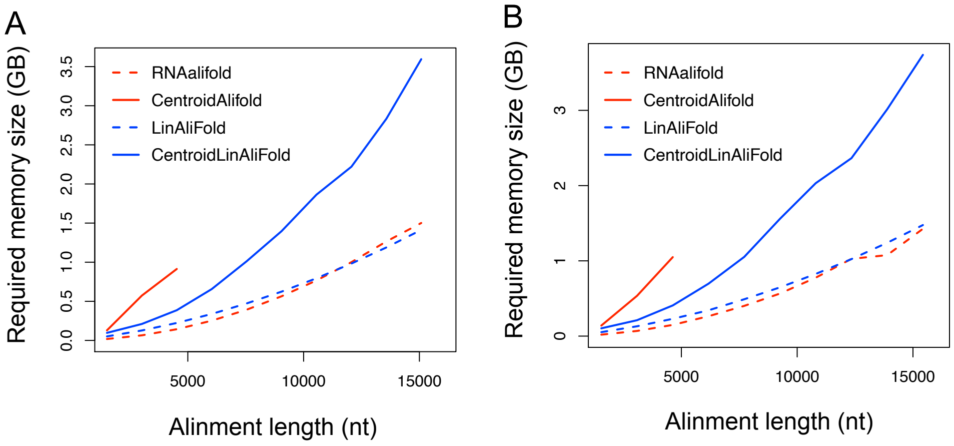


Fig. S3 The required memory size was evaluated for a sequence size of (A) 5 and (B) 10. The x- and y-axis represent the alignment lengths (nt) and required memory size (GB), respectively. The lines represent RNAalifold (red dashed), CentroidAlifold (red solid), LinAliFold (blue dashed), and CentroidLinAliFold (blue solid). The required memory size was not measured when computational time exceeded 2 hours.

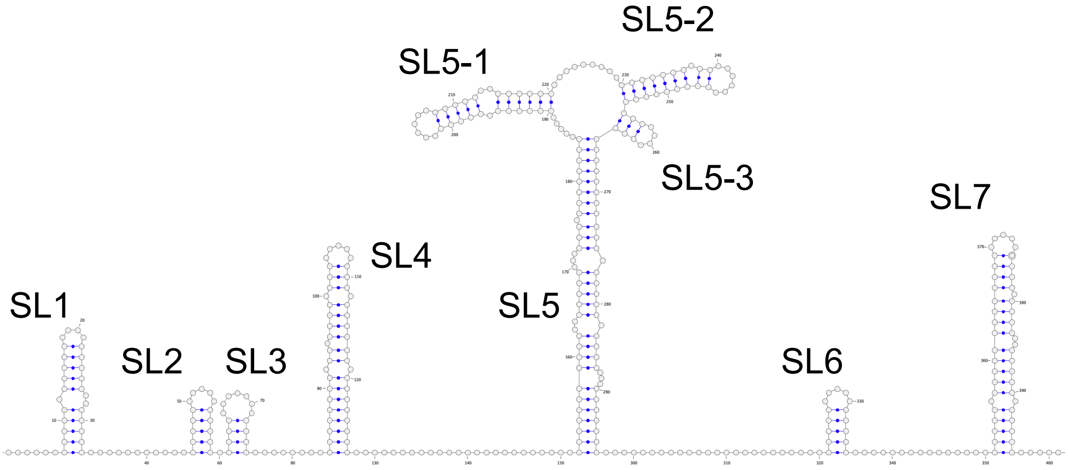


Fig. S4 A predicted RNA consensus secondary structure of coronavirus 5′-UTR by LinAliFold. Graphical view was created using VARNA.

#### Supplementary Tables

Table S1 The PPV results of prediction accuracy for each RNA family when the sequence size was 5

| RNA Family | RNAfold | CentroidFold | RNAalifold | LinAliFold | CentroidAlifold | CentroidLinAliFold |
|---|---|---|---|---|---|---|
| 5s rRNA | 0.757 | 0.742 | 0.902 | 0.886 | **0.914** | **0.914** |
| 16S rRNA | 0.444 | 0.614 | 0.686 | 0.668 | **0.877** | 0.868 |
| group I intron | 0.766 | 0.843 | 0.818 | 0.816 | **0.836** | 0.835 |
| RNaseP | 0.562 | 0.712 | 0.685 | 0.684 | **0.782** | 0.765 |
| SRP RNA | 0.633 | 0.640 | 0.684 | **0.706** | 0.674 | 0.671 |
| telomerase RNA | 0.425 | 0.503 | 0.604 | 0.613 | 0.705 | **0.714** |
| tmRNA | 0.527 | 0.637 | **0.704** | 0.659 | 0.693 | **0.704** |
| tRNA | 0.825 | 0.832 | 0.973 | 0.973 | 0.973 | **0.977** |
| average | 0.618 | 0.691 | 0.757 | 0.751 | **0.807** | 0.806 |

The bold values are the highest scores in each row.

Table S2 The SEN results of prediction accuracy for each RNA family when the sequence size was 5

| RNA Family | RNAfold | CentroidFold | RNAalifold | LinAliFold | CentroidAlifold | CentroidLinAliFold |
|---|---|---|---|---|---|---|
| 5s rRNA | 0.760 | 0.661 | **0.890** | 0.854 | 0.868 | 0.869 |
| 16S rRNA | 0.448 | 0.445 | 0.661 | 0.643 | 0.640 | **0.668** |
| group I intron | **0.704** | 0.686 | 0.472 | 0.488 | 0.520 | 0.491 |
| RNaseP | **0.609** | 0.556 | 0.468 | 0.468 | 0.482 | 0.500 |
| SRP RNA | **0.691** | 0.652 | 0.532 | 0.552 | 0.559 | 0.566 |
| telomerase RNA | 0.561 | 0.567 | 0.572 | 0.582 | 0.579 | **0.588** |
| tmRNA | **0.542** | 0.529 | 0.328 | 0.304 | 0.389 | 0.368 |
| tRNA | 0.838 | 0.774 | **0.970** | **0.970** | **0.970** | 0.937 |
| average | **0.644** | 0.609 | 0.612 | 0.608 | 0.626 | 0.623 |

The bold values are the highest scores in each row.

Table S3 The MCC results of prediction accuracy for each RNA family when the sequence size was 5

| RNA Family | RNAfold | CentroidFold | RNAalifold | LinAliFold | CentroidAlifold | CentroidLinAliFold |
|---|---|---|---|---|---|---|
| 5s rRNA | 0.757 | 0.697 | **0.895** | 0.869 | 0.890 | 0.890 |
| 16S rRNA | 0.445 | 0.522 | 0.673 | 0.655 | 0.748 | **0.760** |
| group I intron | 0.734 | **0.759** | 0.615 | 0.626 | 0.657 | 0.637 |
| RNaseP | 0.584 | **0.627** | 0.563 | 0.562 | 0.610 | 0.616 |
| SRP RNA | **0.658** | 0.641 | 0.584 | 0.606 | 0.605 | 0.608 |
| telomerase RNA | 0.487 | 0.533 | 0.586 | 0.595 | 0.635 | **0.645** |
| tmRNA | 0.533 | **0.577** | 0.467 | 0.436 | 0.514 | 0.502 |
| tRNA | 0.830 | 0.799 | **0.971** | **0.971** | **0.971** | 0.955 |
| average | 0.628 | 0.644 | 0.669 | 0.665 | **0.704** | 0.702 |

The bold values are the highest scores in each row.

Table S4 The PPV results of prediction accuracy for each RNA family when the sequence size was 10

| RNA Family | RNAfold | CentroidFold | RNAalifold | LinAliFold | CentroidAlifold | CentroidLinAliFold |
|---|---|---|---|---|---|---|
| 5s rRNA | 0.735 | 0.732 | 0.934 | 0.913 | **0.940** | 0.919 |
| 16S rRNA | 0.431 | 0.584 | 0.733 | 0.660 | **0.893** | 0.836 |
| group I intron | 0.732 | 0.830 | 0.837 | 0.853 | **0.860** | 0.843 |
| RNaseP | 0.555 | 0.727 | 0.711 | 0.697 | **0.795** | 0.787 |
| SRP RNA | 0.624 | 0.647 | **0.784** | 0.728 | 0.729 | 0.726 |
| telomerase RNA | 0.407 | 0.479 | 0.649 | 0.659 | **0.776** | 0.701 |
| tmRNA | 0.490 | 0.599 | 0.714 | **0.723** | 0.710 | 0.706 |
| tRNA | 0.785 | 0.791 | **0.964** | **0.964** | **0.964** | **0.964** |
| average | 0.595 | 0.674 | 0.791 | 0.775 | **0.833** | 0.810 |

The bold values are the highest scores in each row.

Table S5 The SEN results of prediction accuracy for each RNA family when the sequence size was 10

| RNA Family | RNAfold | CentroidFold | RNAalifold | LinAliFold | CentroidAlifold | CentroidLinAliFold |
|---|---|---|---|---|---|---|
| 5s rRNA | 0.733 | 0.660 | 0.870 | **0.889** | 0.847 | 0.867 |
| 16S rRNA | 0.441 | 0.440 | **0.675** | 0.623 | 0.594 | 0.595 |
| group I intron | **0.670** | **0.670** | 0.481 | 0.497 | 0.476 | 0.498 |
| RNaseP | **0.598** | 0.567 | 0.506 | 0.529 | 0.536 | 0.538 |
| SRP RNA | **0.685** | 0.676 | 0.596 | 0.625 | 0.606 | 0.621 |
| telomerase RNA | 0.532 | 0.537 | 0.400 | **0.552** | 0.454 | **0.552** |
| tmRNA | **0.516** | 0.501 | 0.247 | 0.291 | 0.332 | 0.336 |
| tRNA | 0.798 | 0.737 | **0.959** | **0.959** | **0.959** | **0.959** |
| average | **0.622** | 0.599 | 0.592 | 0.621 | 0.600 | 0.621 |

The bold values are the highest scores in each row.

Table S6 The MCC results of prediction accuracy for each RNA family when the sequence size was 10

| RNA Family | RNAfold | CentroidFold | RNAalifold | LinAliFold | CentroidAlifold | CentroidLinAliFold |
|---|---|---|---|---|---|---|
| 5s rRNA | 0.732 | 0.692 | **0.900** | **0.900** | 0.891 | 0.892 |
| 16S rRNA | 0.435 | 0.506 | 0.703 | 0.640 | **0.726** | 0.704 |
| group I intron | 0.700 | **0.744** | 0.631 | 0.647 | 0.637 | 0.645 |
| RNaseP | 0.575 | 0.640 | 0.595 | 0.602 | **0.649** | 0.646 |
| SRP RNA | 0.650 | 0.657 | **0.671** | 0.669 | 0.659 | 0.665 |
| telomerase RNA | 0.464 | 0.506 | 0.505 | 0.599 | 0.590 | **0.617** |
| tmRNA | 0.501 | **0.544** | 0.416 | 0.454 | 0.477 | 0.478 |
| tRNA | 0.789 | 0.759 | **0.961** | **0.961** | **0.961** | **0.961** |
| average | 0.606 | 0.631 | 0.673 | 0.684 | 0.699 | **0.701** |

The bold values are the highest scores in each row.

Table S7 The PPV results of prediction accuracy for each RNA family when the sequence size was 20

| RNA Family | RNAfold | CentroidFold | RNAalifold | LinAliFold | CentroidAlifold | CentroidLinAliFold |
|---|---|---|---|---|---|---|
| 5s rRNA | 0.730 | 0.716 | **0.940** | 0.918 | **0.940** | 0.925 |
| 16S rRNA | 0.431 | 0.607 | 0.726 | 0.677 | **0.892** | 0.835 |
| group I intron | 0.738 | 0.836 | 0.833 | 0.820 | **0.845** | 0.823 |
| RNaseP | 0.552 | 0.743 | 0.748 | 0.689 | 0.759 | **0.768** |
| SRP RNA | 0.616 | 0.631 | 0.770 | 0.748 | **0.775** | 0.748 |
| telomerase RNA | 0.395 | 0.461 | 0.726 | 0.632 | **0.800** | 0.715 |
| tmRNA | 0.504 | 0.607 | 0.744 | 0.686 | **0.749** | 0.665 |
| tRNA | 0.804 | 0.815 | **0.978** | 0.972 | **0.978** | 0.972 |
| average | 0.596 | 0.677 | 0.808 | 0.768 | **0.842** | 0.806 |

The bold values are the highest scores in each row.

Table S8 The SEN results of prediction accuracy for each RNA family when the sequence size was 20

| RNA Family | RNAfold | CentroidFold | RNAalifold | LinAliFold | CentroidAlifold | CentroidLinAliFold |
|---|---|---|---|---|---|---|
| 5s rRNA | 0.732 | 0.646 | 0.836 | 0.879 | 0.836 | **0.884** |
| 16S rRNA | 0.438 | 0.449 | **0.645** | 0.636 | 0.607 | 0.622 |
| group I intron | **0.680** | 0.679 | 0.433 | 0.492 | 0.467 | 0.508 |
| RNaseP | **0.588** | 0.568 | 0.513 | 0.522 | 0.536 | 0.560 |
| SRP RNA | **0.680** | 0.659 | 0.489 | 0.586 | 0.495 | 0.595 |
| telomerase RNA | 0.528 | **0.532** | 0.263 | 0.491 | 0.385 | 0.465 |
| tmRNA | **0.524** | 0.500 | 0.209 | 0.340 | 0.291 | 0.390 |
| tRNA | 0.819 | 0.763 | 0.948 | **0.961** | 0.948 | **0.961** |
| average | **0.624** | 0.600 | 0.542 | 0.614 | 0.571 | 0.623 |

The bold values are the highest scores in each row.

Table S9 The dependencies of the computational speeds (s) on the sequence size

| software | concatenation size | sequence size 5 | 10 | 20 |
|---|---|---|---|---|
| LinAliFold | 1 | 4.77 | 5.67 | 7.57 |
| | 5 | 29.19 | 33.06 | 44.39 |
| | 10 | 62.60 | 80.04 | 143.58 |
| CentroidLinAliFold | 1 | 31.03 | 43.73 | 79.09 |
| | 5 | 295.29 | 407.71 | 653.50 |
| | 10 | 954.09 | 1256.32 | 1892.06 |

Table S10 The dependencies of the required memory size (GB) on the sequence size

| software | concatenation size | sequence size 5 | 10 | 20 |
|---|---|---|---|---|
| LinAliFold | 1 | 0.05 | 0.05 | 0.05 |
| | 5 | 0.47 | 0.49 | 0.51 |
| | 10 | 1.41 | 1.48 | 1.17 |
| CentroidLinAliFold | 1 | 0.10 | 0.10 | 0.11 |
| | 5 | 1.01 | 1.05 | 1.11 |
| | 10 | 3.60 | 3.74 | 3.79 |

Table S11 The dependencies of the MCC results on the covariance model when the sequence size was 20

| | RNAalifold | | LinAliFold | | CentroidLinAlifold | |
|---|---|---|---|---|---|---|
| RNA Family | simple | RIBOSUM | simple | RIBOSUM | simple | RIBOSUM |
| 5s rRNA | 0.885 | 0.919 | 0.898 | 0.918 | 0.903 | 0.932 |
| 16S rRNA | 0.684 | 0.675 | 0.656 | 0.658 | 0.720 | 0.735 |
| group I intron | 0.598 | 0.621 | 0.632 | 0.626 | 0.644 | 0.642 |
| RNaseP | 0.616 | 0.685 | 0.596 | 0.677 | 0.654 | 0.705 |
| SRP RNA | 0.596 | 0.631 | 0.645 | 0.634 | 0.649 | 0.645 |
| telomerase RNA | 0.436 | 0.573 | 0.555 | 0.567 | 0.572 | 0.600 |
| tmRNA | 0.382 | 0.510 | 0.475 | 0.522 | 0.505 | 0.525 |
| tRNA | 0.962 | 0.966 | 0.966 | 0.966 | 0.966 | 0.966 |
| average | 0.645 | 0.698 | 0.678 | 0.696 | 0.702 | 0.719 |

Table S12 The dependencies of the MCC results of LinAliFold on the beam size when the sequence size was 20

| RNA Family | $b$=10 | $b$=30 | $b$=50 | $b$=100 | $b$=200 | $b$=500 |
|---|---|---|---|---|---|---|
| 5s rRNA | 0.881 | 0.898 | 0.898 | 0.898 | 0.898 | 0.898 |
| 16S rRNA | 0.457 | 0.613 | 0.651 | 0.656 | 0.714 | 0.738 |
| group I intron | 0.362 | 0.580 | 0.609 | 0.632 | 0.632 | 0.632 |
| RNaseP | 0.514 | 0.596 | 0.596 | 0.596 | 0.596 | 0.596 |
| SRP RNA | 0.645 | 0.645 | 0.645 | 0.645 | 0.645 | 0.645 |
| telomerase RNA | 0.285 | 0.459 | 0.490 | 0.555 | 0.563 | 0.563 |
| tmRNA | 0.403 | 0.467 | 0.475 | 0.475 | 0.475 | 0.475 |
| tRNA | 0.936 | 0.966 | 0.966 | 0.966 | 0.966 | 0.966 |
| average | 0.560 | 0.653 | 0.666 | 0.678 | 0.686 | 0.689 |

Table S13 The dependencies of the MCC results of CentroidLinAliFold on the beam size when the sequence size was 20

| RNA Family | $b$=10 | $b$=30 | $b$=50 | $b$=100 | $b$=200 | $b$=500 |
|---|---|---|---|---|---|---|
| 5s rRNA | 0.900 | 0.903 | 0.903 | 0.903 | 0.903 | 0.903 |
| 16S rRNA | 0.571 | 0.666 | 0.722 | 0.720 | 0.771 | 0.768 |
| group I intron | 0.518 | 0.622 | 0.640 | 0.644 | 0.644 | 0.644 |
| RNaseP | 0.583 | 0.652 | 0.654 | 0.654 | 0.653 | 0.653 |
| SRP RNA | 0.644 | 0.649 | 0.649 | 0.649 | 0.649 | 0.649 |
| telomerase RNA | 0.470 | 0.538 | 0.557 | 0.572 | 0.574 | 0.575 |
| tmRNA | 0.417 | 0.511 | 0.504 | 0.505 | 0.511 | 0.511 |
| tRNA | 0.944 | 0.966 | 0.966 | 0.966 | 0.966 | 0.966 |
| average | 0.631 | 0.688 | 0.699 | 0.702 | 0.709 | 0.709 |

Table S14 The dependencies of the computational speeds (s) on the beam size when the sequence size was 20

| | | beam size | | | | | |
|---|---|---|---|---|---|---|---|
| software | concatenation size | 10 | 30 | 50 | 100 | 200 | 500 |
| LinAliFold | 1 | 1.11 | 2.70 | 4.24 | 7.58 | 12.66 | 20.13 |
| | 5 | 5.75 | 15.03 | 22.95 | 58.77 | 91.19 | 218.62 |
| | 10 | 15.73 | 38.96 | 63.16 | 143.58 | 261.05 | 710.48 |
| CentroidLinAliFold | 1 | 11.396 | 26.21 | 44.61 | 79.09 | 153.56 | 304.20 |
| | 5 | 156.24 | 247.67 | 360.23 | 653.50 | 1569.04 | 5728.0 |
| | 10 | 709.70 | 941.82 | 1171.00 | 1892.06 | 4245.25 | - |

'-' represents that the computational time was longer than 2 hours.

Table S15 The dependencies of the required memory size (GB) on the beam size when the sequence size was 20

| software | concatenation size | beam size 10 | 30 | 50 | 100 | 200 | 500 |
|---|---|---|---|---|---|---|---|
| LinAliFold | 1 | 0.02 | 0.03 | 0.04 | 0.05 | 0.08 | 0.14 |
| | 5 | 0.29 | 0.34 | 0.40 | 0.51 | 0.69 | 0.87 |
| | 10 | 1.07 | 1.17 | 1.29 | 1.17 | 1.76 | 2.10 |
| CentroidLinAliFold | 1 | 0.07 | 0.08 | 0.08 | 0.11 | 0.14 | 0.16 |
| | 5 | 1.07 | 1.07 | 1.09 | 1.11 | 1.16 | 2.03 |
| | 10 | 3.43 | 3.37 | 3.70 | 3.79 | 3.82 | - |

'-' represents that the computational time was longer than 2 hours.
